# Cardiovascular and vasomotor pulsations in the brain and periphery during awake and NREM sleep in a multimodal fMRI study

**DOI:** 10.1101/2024.04.02.587690

**Authors:** Johanna Tuunanen, Heta Helakari, Niko Huotari, Tommi Väyrynen, Matti Järvelä, Janne Kananen, Annastiina Kivipää, Lauri Raitamaa, Seyed-Mohsen Ebrahimi, Mika Kallio, Johanna Piispala, Vesa Kiviniemi, Vesa Korhonen

## Abstract

The glymphatic brain clearance mechanism convects brain cerebrospinal fluid driven by physiological pulsations such as cardiovascular and very low-frequency (VLF < 0.1 Hz) vasomotor waves. Presently, ultrafast functional magnetic resonance imaging (fMRI) facilitates the measurement of these signals from both venous and arterial compartments. In this study, we compared the interaction of these two pulsations in awake and sleep using fMRI and peripheral fingertip photoplethysmography in both arterial and venous signals in ten subjects (5 female). Sleep increased the power of brain cardiovascular pulsations, decreased peripheral pulsation and desynchronized them. Vasomotor waves, however, increase in both power and synchronicity in brain and peripheral signals during sleep. Peculiarly, vasomotor lag reversed in sleep within the default mode network vs. peripheral signal. Finally, sleep synchronized cerebral arterial vasomotion measured with cardiovascular hemodynamic envelope (CHe) vs. venous blood oxygenation level dependent (BOLD) signals in parasagittal brain tissue. These changes in power and pulsation synchrony may reflect differential changes in vascular control between the periphery and brain vasculature, while the increased synchrony of arterial and venous compartments may reflect increased convection of neurofluids in parasagittal areas in sleep.

**Statement of Significance:** This study shows that while sleep attenuated the cardiovascular synchrony and powers of pulsatility between the periphery and brain, it also increased brain tissue synchrony of venous and arterial vasomotor waves, specifically in the parasagittal regions. The study also shows that vasomotor waves increased in the human brain and the periphery during NREM sleep. Thus, sleep induces a whole-body vasomotor synchronization where the initiation of peripheral vasomotor waves is preceded within the default mode network area. Based on these results, we suggest that the synchronization of vasomotor waves may be a significant contributor to the enhancement of glymphatic fluid exchange in the human brain during sleep.

## Introduction

Angelo Mosso demonstrated in 1880 that cerebral blood flow and cardiovascular brain pulsations increased due to increased brain activity in a patient with a cranial defect^1^. Similar increases in arterial pulsatility can now be non-invasively detected using fast functional magnetic resonance imaging (fMRI) measurements that can capture the whole brain at 10 Hz sampling frequency, thus allowing sufficient temporal resolution of each arterial impulse, without any aliasing^2,3^. The dissociation of cerebral arterial and venous signals is possible due to their differing i.e. spin phase vs. susceptibility magnetic resonance (MR) signal generation mechanisms^4–7^. Setting a cardiovascular hemodynamic envelope (CHe)^8^ over each arterial impulse peak allows the detection of the local arterial dilations, which closely follow brain activations as directed by neurovascular coupling. The CHe signal was shown to precede the venous blood oxygenation level dependent (BOLD) signal by approximately 1.3 seconds in visual cortical areas^8^. Also, breath-hold stimulation caused changes in the respiratory brain stem CHe signal^9^. Previously, the coupling of peripheral vascular waves with BOLD signal changes in the brain has shown to be consistent^10,11^.

During sleep, the amplitude of the cardiovascular impulses and of the three known physiological brain pulsations, namely very low-frequency (VLF < 0.1 Hz), respiration, and cardiac waves, increase in power along with the transition of brain electrophysiological signals towards slow delta electroencephalography (EEG) oscillations^12^. The increased amplitude in these various pulsations drives blood and cerebrospinal fluid (CSF) flow to maintain brain homeostasis. In particular, the pulsations drive the transfer of solutes from periarterial CSF spaces via interstitial fluid (ISF) spaces into perivenous conduits^13–16^, in what has come to be known as the glymphatic clearance pathway. At least in sub-arachnoid periarterial spaces, the main driver of hydrodynamic exchange is the cardiovascular pulsations^13,14,17^. Moreover, the slow (< 0.1 Hz) vasomotor waves of smooth muscle tone in arterial walls of mice impact the brain solute convection along the blood vessel basement membranes^18^. The basement membrane, which extends into the interstitial space, can serve as the exchange route for solute trafficking both in periarterial and perivenous spaces^19^.

Imaging of the pulse amplitudes simultaneously both in arterial and venous compartments allows a new perspective into understanding the physiology of cerebral blood flow and solute transport. Synchrony between the pulses of these two compartments could reflect the state of the conduits linking the arterial and venous sides; Increased synchrony could be related to increased flow conductance between the compartments, whereby arterial flow changes would increasingly drive venous clearance. Furthermore, larger arterial impulse amplitudes correlate with better convection of solutes, which is characterized by less backward flow in pial periarterial spaces^14^.

In this study, we used a fast fMRI magnetic resonance encephalography (MREG) sequence to investigate the changes in power and the pulsatile synchrony between arterial and venous brain compartments in the transition to sleep in healthy volunteers. We thereby tested the hypothesis that the increased power of the pulsations during sleep would synchronize the interaction between the venous and arterial spaces as compared to the awake state. At first, we confirmed the accuracy of MREG for detecting pulsations against the peripheral fingertip photoplethysmography (PPG). We then compared EEG-verified sleep and awake data to assess how sleep alters cardiovascular pulsatility (MREG_CARD_), arterial vasomotor activity waves (MREG_CHe_), and classical venous BOLD signal (MREG_BOLD_).

## Materials and methods

### Subjects

The analysis was performed on ten healthy volunteers (age in years: 25.5 ± 3.4, 5 females). The study was approved by the Regional Ethics Committee of the Northern Ostrobothnia Hospital District and was performed in Oulu, Finland. Written informed consent was obtained from all participants according to the Declaration of Helsinki. All subjects were healthy and met the following inclusion criteria: no continuous medication, no neurological nor cardio-respiratory diseases, non-smokers, and no pregnancy. The subjects were instructed not to consume caffeine during the four hours before the awake scan session and eight hours before the sleep scan session. Participants agreed to abstain from alcohol consumption 12 hours before the scans.

### Data acquisition

The subjects participated in two separate measurement sessions three days apart, with an awake scan session in the afternoon and a sleep scan session in the early morning after 24 hours of sleep deprivation. Participants wore an Ōura Ring (ouraring.com) for at least 24 hours preceding both imaging sessions in a Siemens MAGNETOM 3T Skyra scanner (Siemens Healthineers AG, Erlangen, Germany) equipped with a 32-channel head-coil. We used a fast fMRI imaging sequence MREG, consisting of a 3D single shot stack of spiral sequences that under-samples k-space, which allows sampling of physiological pulsations at 10 Hz^20^. The following scanning parameters were used for MREG: repetition time (TR=100 ms), echo time (TE=36 ms), field of view (FOV=192 mm), flip angle (FA=5°), and 3 mm isotropic cubic voxel. MREG scan images were reconstructed using L2-Tikhonov regularization with lambda 0.1, with the latter regularization parameter having been determined by the L-curve method using a MATLAB recon-tool provided by the sequence developers^21^. Anatomical 3D structural T1 MPRAGE (TR=1900 ms, TE=2.49 ms, FOV=240 mm, FA=9° and 0.9 mm slice thickness) images were used to register MREG datasets into Montreal Neurological Institute (MNI) space.

During the awake scan session, one 10-minute resting state MREG sequence was taken with eyes open and gaze fixated on a cross projected on a monitor. During the sleep measurement session three days later, two 10-minute MREG scans were first taken. Before the actual fMRI scan, sleep-deprived participants were instructed to lie down on the scanner bed, close their eyes, and attempt to sleep, accompanied by the dimming of lights in the MRI room. An anatomical T1-weighted scan was performed at the end of both sessions. Part of the research pipeline is shown in Fig. 1, with greater details presented in Helakari et al. (2022)^12^.

**Figure 1.**
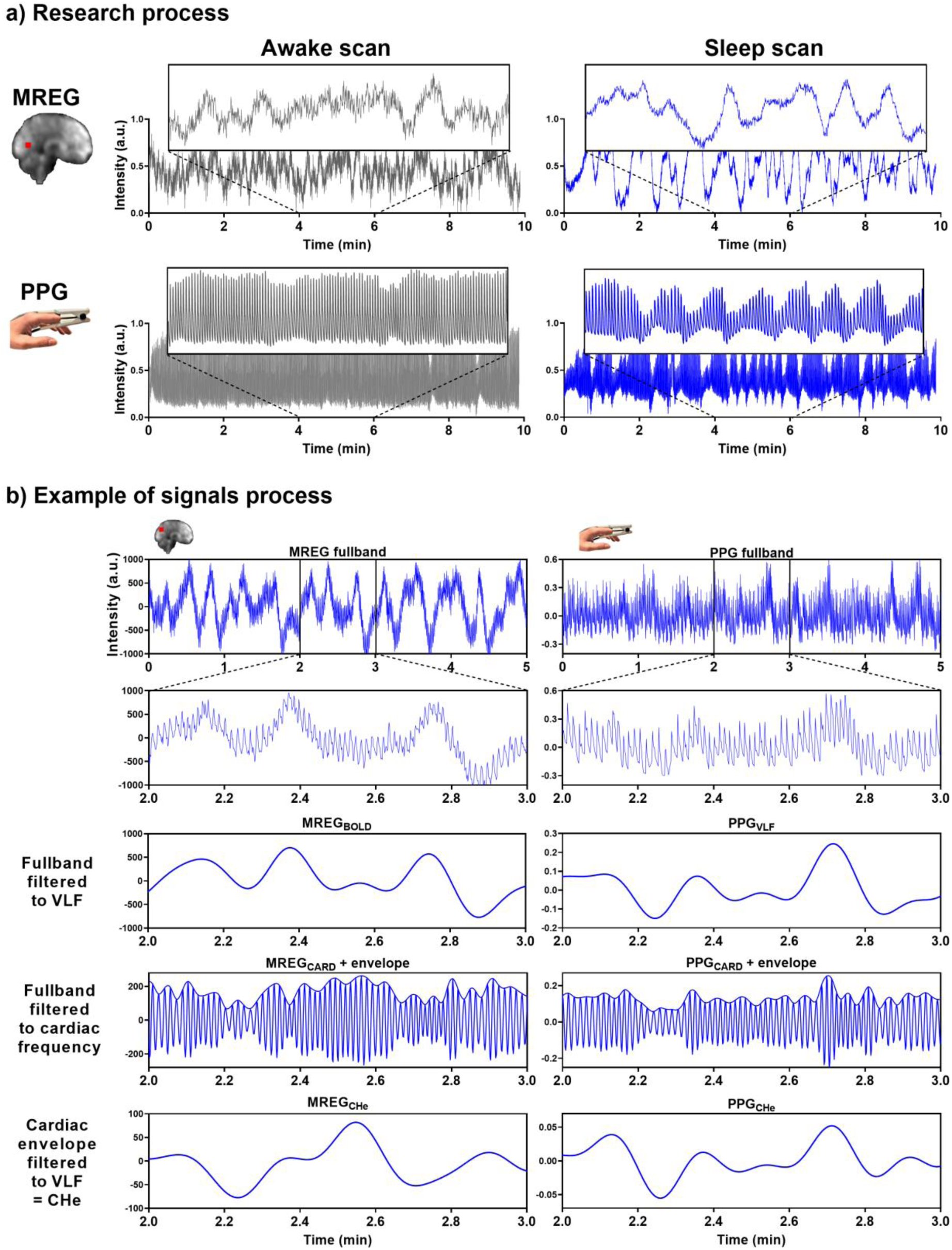
Representative signals for awake and NREM sleep. a) During the awake scan (10 minutes) eyes were open and during the sleep scans (2×10 minutes) subjects were allowed to fall asleep. MREG and PPG signals are from an awake (5 % sleep, EEG scored) subject and from a sleeping (85 % sleep, EEG scored) subject. Example signals are normalized. b) In the analysis we used 5-minute epoch signals. Examples of MREG and PPG signal processing in a sleeping (30 % NREM N1 and 70 % NREM N2 EEG scored sleep) subject.

Fingertip peripheral PPG, electrocardiography (ECG), and end-tidal-CO_2_ (EtCO_2_) were measured in synchrony using a 3-T MRI-compatible anesthesia monitor (Datex-Ohmeda S/5 Collect software), as described previously^22^. Cuff-based blood pressure of each subject was also measured sitting and supine before both scanning sessions. The PPG probe was placed on the left index finger, with data acquisition at 300 Hz. Right index fingertip PPG data were collected using the MR scanner for verification purposes.

### Pre-processing

Preprocess steps and analyses were performed using MATLAB (R2020b, The MathWorks, Natick, MA), Functional MRI of the Brain Software Library (FSL; Brain Extraction Tool (BET), version 5.09)^23,24^ and Analysis of Functional NeuroImages (AFNI; version 2)^25^. As a part of quality control, we visually inspected the PPG data. We identified and corrected high peaks caused by motion artifacts by despike, allowing the use of the mean value for adjacent PPG amplitudes during analysis. Raw PPG data were used to calculate cardiorespiratory parameters and frequency ranges, and then down-sampled to 10 Hz using MATLAB for analysis of fast Fourier transform (FFT) power and cross-correlation with MREG data.

FSL BET was used for brain extraction with neck and bias-field correction from structural 3D MPRAGE volumes^24^. MREG datasets passed through a typical FSL preprocessing pipeline^23^, with high-pass filtering at a cutoff of 0.008 Hz. Datasets were spatially smoothed with 5 mm Full Width at Half Maximum Gaussian kernel. AFNI *3dDespike* was used to remove spikes from the datasets caused by head movements. Head motion was calculated with FSL 5.08 MCFLIRT (Functional MRI of the Brain Linear Image Registration Tool)^26^. MCFLIRT relative or absolute mean displacement values (mm) did not differ significantly between awake and sleep states (p_rel_ = 0.46, p_abs_ = 0.07).

To get the maximum amount of awake signal from awake scans and likewise, NREM N2 sleep from sleep scans, PPG and MREG signals were sectioned into continuous simultaneous 5-minute segments (3000-time points) using MATLAB and *fslroi* (Fig. 1b). The very low-frequency (VLF) range of 0.008-0.1 Hz was chosen to get maximum possible coverage of the low frequencies without crosstalk with respiratory frequencies. Cardiac frequency ranges were obtained from individual PPG spectrums without respiratory aliasing, harmonics, or noise for correlation analysis. After analysis, the data were registered into MNI space at 3 mm resolution to enable comparable analysis between awake and sleep datasets.

EEG was recorded using the Electrical Geodesics MR-compatible GES 400 system (Magstim), with a 256-channel high-density scalp net. Electrode impedances were < 50 kΩ and the sampling rate was 1 kHz. Signal quality was tested outside the scanner room by recording 30-second epochs of EEG with eyes open and eyes closed.

### Cardiorespiratory analysis

PPG and EtCO_2_ signals from an anesthesia monitor were used to verify group cardiorespiratory parameters based on the individual ranges of each subject. Heart rate (HR) from the PPG signal and respiratory rate (RR) from the EtCO2 signal were determined using the MATLAB *findpeaks* function with subject-specific input parameters. RR of one subject was determined from MREG data^27,28^ due to invalid respiration data. Heart rate variability (HRV) was calculated from the PPG signal to determine the root mean square of successive differences between normal heartbeats^29^ for awake and sleep scans. HR, HRV, and RR were calculated from one 5-minute awake scan and one 5-minute sleep scan, which were selected for the analysis.

### Power analysis of PPG and global MREG

The down-sampled 5-minute PPG signals were normalized using Z-score normalization^30^ and MATLAB *fft* function was used to calculate the respective FFT power spectra. Z-score normalization ensured that the amplitudes of the PPGs were consistent. Global FFT power density maps were calculated using AFNI *3dPeriodogram* function for 5 minutes of full band (0.008 – 5 Hz) MREG data, followed by application of the *fslmeants* function to calculate the average FFT spectrum of all brain voxels. The VLF and individual cardiac FFT power were calculated using AFNI *3dTstat*. The VLF and cardiac band powers and mean (+ standard deviation) power spectra were visualized using GraphPad Prism 9 software. The most prominent individual cardiac peaks were shifted to the group mean frequency of both awake and sleep scans, and the mean power spectrum was calculated for each condition.

### Cross-correlation analysis

The voxel-wise positive maximum cross-correlation coefficients (CC_MAX_) between the pre-processed PPG and MREG signals in awake and sleep states were calculated using the MATLAB *xcorr* function with sequence normalization *‘coeff’* and the corresponding time delays yielding the maximum correlation. We restricted the search range of time delays to be within ±10 s, as the total transit time of blood through the head is less than 10 seconds in healthy subjects^31^. Cross-correlation was performed between the following 5-minute signals:

1. **Fingertip cardiac pulsatility to cerebral cardiac pulsatility correlation.** Cardiac signals reflected the resultant arterial pulsation. Here, the PPG signal was first analyzed with FFT and was then band-pass filtered to each subject’s individual cardiac frequency band^28^ using MATLAB. First, the raw MREG data were multiplied by −1 to raise the R-peak. Next, using the individual frequency information, the full-band MREG data were band-pass filtered to each subject’s cardiac range using the AFNI *3dTproject* to obtain signal without noise, respiratory aliasing, or harmonic cardiac waves. These cardiac-filtered signals are henceforth designated as PPG_CARD_ and MREG_CARD_.
2. **Cerebral arterial vasomotor wave vs. venous BOLD signal correlation.** The brain MREG signal contains two forms of very low-frequency (VLF, 0.008-0.1 Hz) vascular waves in addition to other physiological signals; there is an active *arterial* vasomotor tone wave, which is followed some 1.3 sec later by a passive *venous* wave resulting from downstream ballooning due to blood volume and oxygenation changes^8^. Since these two waves have a different MR contrast origin (arterial spin phase vs. venous susceptibility), they can be separated from fast MREG data with the following procedure: Cerebral MREG_CHe_ reflecting *arterial vasomotor tone waves* can be obtained from peak-to-peak cardiovascular impulse amplitudes using a cardiovascular hemodynamic envelope (MREG_CHe,_ c.f. Huotari et al., 2022). The MREG_CHe_ were obtained by setting an envelope over the individually verified cardiac frequency MREG_CARD_ signal peaks with MATLAB *findpeaks*, *spline*, and *ppval* functions. The classical BOLD baseline signal reflecting venous blood volume and (de)oxygenation level fluctuations were obtained, importantly also without aliased physiological noise, harmonics, and modulations, by bandpass filtering the full band MREG data to the VLF range using AFNI *3dTproject*; we henceforth designate this as MREG_BOLD_. To analyze similar physiological phenomena, the MREG_CHe_ data were also bandpassed to the *identical (0.008-0.1 Hz)* VLF range. Finally, the MREG_CHe_ vs. MREG_BOLD_ were cross-correlated on a voxel-by-voxel basis.
3. **Fingertip VLF blood volume waves to MREG_BOLD_ correlation.** Fingertip PPG signals reflect blood volume changes^32,33^. The PPG signal was bandpass filtered to the VLF band using MATLAB, which we designate as the peripheral PPG_VLF_; this was also cross-correlated to MREG_BOLD_.
4. **Fingertip vasomotor arterial wave to MREG_BOLD_ correlation**. Peripheral vasomotor tone (a.k.a. arterial amplitude) waves were also estimated from the fingertip PPG signal using the CHe technique as described above in item 2. The PPG signal was bandpass-filtered to the individual cardiac range using MATLAB to obtain a signal uncontaminated with other physiological signals. An envelope was set above the cardiac peaks for extraction of the PPG_CHe_ similar as above. Finally, the envelopes were bandpass filtered to the VLF range for correlation analysis.

### Preprocessing and analysis of EEG data

EEG recordings were preprocessed using the Brain Vision Analyzer (Version 2.1; Brain Products) after converting to a compatible format via BESA Research (Version 7.0). Gradient artifacts due to static and dynamic magnetic fields during the MRI data acquisition, along with the ballistocardiograph (BCG) artifacts, were corrected using the average artifact subtraction method^34,35^. The data was checked to ensure the absence of gradient and BCG artifacts.

After preprocessing, EEG data were visualized according to the 10-20 system for sleep state scoring. The EEG data were scored in 30 s epochs by two experienced specialists in clinical neurophysiology following the American Academy of Sleep Medicine (AASM) 2017 guidelines, and the final sleep state scoring was obtained by consensus of both specialists. Using established criteria, EEG epochs were scored as wake, NREM N1 (light sleep), NREM N2 (intermediate sleep with sleep spindles and/or K-complexes), NREM N3 (slow wave sleep), or REM (sleep with rapid eye movements).

### Ōura Ring activity data

The subject sleep/wake status was monitored with the multisensory Ōura Ring sleep tracker (ouraring.com) on the days before each scan, with a detailed analysis of the final 24 hours before scanning. The principles and results of the ring data are presented in Helakari et al. (2022)^12^.

### Cantab tests

At the beginning of scanning sessions, the subjects underwent reaction time and paired-associate learning tests from the Cantab (Cambridge Neuropsychological Test Automated Battery, Cantab, Cambridge, UK) test battery, performed on a tablet computer. The results are shown in Helakari et al. (2022)^12^.

### Statistical analysis

Whole brain voxel-wise comparisons between different awake/sleep states in the same subjects were performed by two-sample t-test using a paired nonparametric threshold-free permutation test (5,000 permutations) implemented in *vlisa_2ndlevel* from LIPSIA^36^. The Shapiro-Wilk test was used to examine the normality of the distribution of the variables. For the sum of PPG FFT power, Oura ring activity data, Cantab test performance, cardiorespiratory signals, and head motion, we calculated statistical differences by two-sided paired Student’s t-test using GraphPad Prism 9. Results are reported as mean ± standard deviation, and p-values below 0.05 are considered significant (p < 0.05: *, p < 0.01: **, p < 0.001: ***, p < 0.0001: ****).

## Results

We investigated cerebral and peripheral cardiovascular pulsatility, as well as vasomotor waves, and examined their relationships during awake and NREM sleep epochs. Subjects were scanned after a monitored normal night sleep and after monitored sleep deprivation (24 h) time. During the awake scan, subjects were awake for 100 % of the five-minute data (Fig. 2a), whereas during the sleep scan, subjects were awake for 4 ± 10 % of the five-minute epochs (Fig. 2a), with 27 ± 31 % N1 sleep and 69 ± 31 % N2 sleep, but no N3 or REM sleep was detectable.

**Figure. 2.**
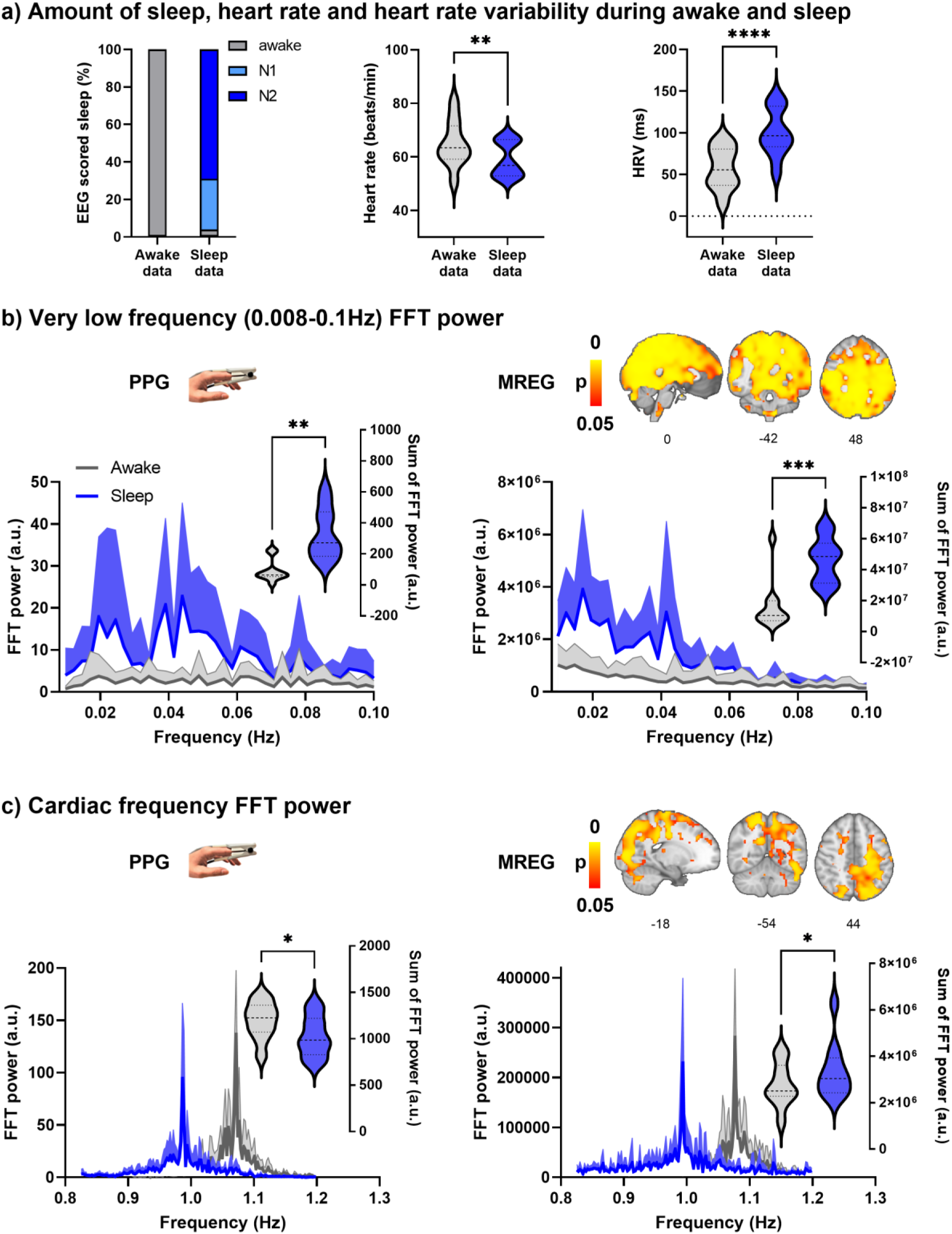
Prevalence of vigilance state in the sleep data and spectral FFT power of fingertip peripheral PPG and MREG recordings during awake and sleep states (n = 10). a) During the sleep recordings, subjects slept (NREM N1-N2 sleep) 96 % of the time. Calculated from PPG, the heart rate (HR, p = 0.0099 **) was significantly lower during sleep, and heart rate variability (HRV, p < 0.0001 ****) significantly higher during sleep compared to awake. b) Group average (+ standard deviation) PPG and global MREG spectra in very low-frequency (VLF, 0.008-0.1 Hz) show that the VLF power was significantly higher during sleep compared to awake in PPG (p = 0.0014, **) and in global MREG data (p = 0.0007, ***). c) The frequency-normalized group mean cardiac frequency spectra showed that cardiac frequency power was significantly lower in fingertip PPG (p = 0.037, *) but higher in global MREG (p = 0.035, *) during sleep compared to awake.

### Cardiorespiratory analysis

Cardiorespiratory parameters were calculated from the fingertip PPG and EtCO_2_ signals from awake and sleep five-minute epochs (Table 1.).

**Table 1.** Cardiorespiratory parameters.

| Outcome | N | Awake data | Sleep data | Mean change | P-value |
| --- | --- | --- | --- | --- | --- |
|  |  | Mean (SD) | Mean (SD) | (95 % CI) |  |
| Heart rate (b/min) | 10 | 65 (9) | 59 (7) | -6 (-10 to -2) | 0.0099<sup>#</sup> |
| Heart rate variability (ms) | 10 | 57 (25) | 100 (29) | 43 (29 to 57) | <0.0001<sup>#</sup> |
| Respiratory rate (b/min) | 10 | 16.3 (3.3) | 15.3 (2.4) | -0.9 (-2.5 to 0.6) | 0.21<sup>#</sup> |
| Cardiac frequency width |  |  |  | 0.12 (0.08 to |  |
| (Hz) | 10 | 0.24 (0.05) | 0.36 (0.04) | 0.17) | 0.0001<sup>#</sup> |

<sup>#</sup>Paired t-test
SD = Standard deviation
CI = Confidence interval

Heart rate (HR) decreased (p = 0.0099, **, Fig. 2a) and heart rate variability (HRV) increased (p < 0.0001, ****, Fig. 2a) significantly during sleep compared to awake. The width of the cardiac spectrum expanded during sleep (p = 0.0001, ***). Respiratory rate (RR) showed no significant differences between sleep and awake recordings. Group mean blood pressure (systole/diastole) was 127/77 mmHg (sitting), 127/69 mmHg (supine) before the awake scan, and 127/75 mmHg (sitting), 125/68 mmHg (supine) before the sleep scan.

### Power analysis of PPG and global MREG

FFT amplitude power was calculated from the fingertip peripheral PPG signal and global MREG signal from awake and sleep data (Fig. 2b-c). The FFT spectra of both the fingertip and the global brain signals show the greatest increase in power during sleep at 0.02 Hz and 0.04 Hz VLF frequencies. FFT power was significantly higher in the VLF band during sleep compared to awake in PPG (p = 0.0014, **) and global MREG (p = 0.0007, ***), much as reported previously by Helakari et al. (2022)^12^. In cardiac frequency, the FFT power was significantly lower in fingertip PPG (p = 0.037 *), but interestingly was higher in the brain global MREG (p = 0.035, *) during sleep compared to awake.

### Beat-to-beat cardiovascular synchrony between periphery and brain

PPG and MREG signals were filtered to the individual cardiac range. Figure 3 illustrates the drastic amplitude modulation and de-synchronization in sleep between the measurement points. As a sign of high temporal accuracy, the MREG_CARD_ signal of the brain gave nearly identical pulsation signals compared to PPG_CARD_, especially in the proximity of major arteries, c.f. Fig. 3c. However, during sleep, the CC_MAX_ between peripheral PPG_CARD_ and brain MREG_CARD_ were significantly lower compared to the awake scan, extending over most parts of the brain and especially in central areas (p < 0.05, Fig. 3d). Group mean lag values ranged between - 1 - +1 sec during awake recordings, and without any significant change in sleep (Fig. 3e). The MREG_CARD_ lead the fingertip lags by approximately 0.3 ± 0.4 seconds in awake scans and 0.4 ± 0.3 seconds in sleep scans.

**Figure 3.**
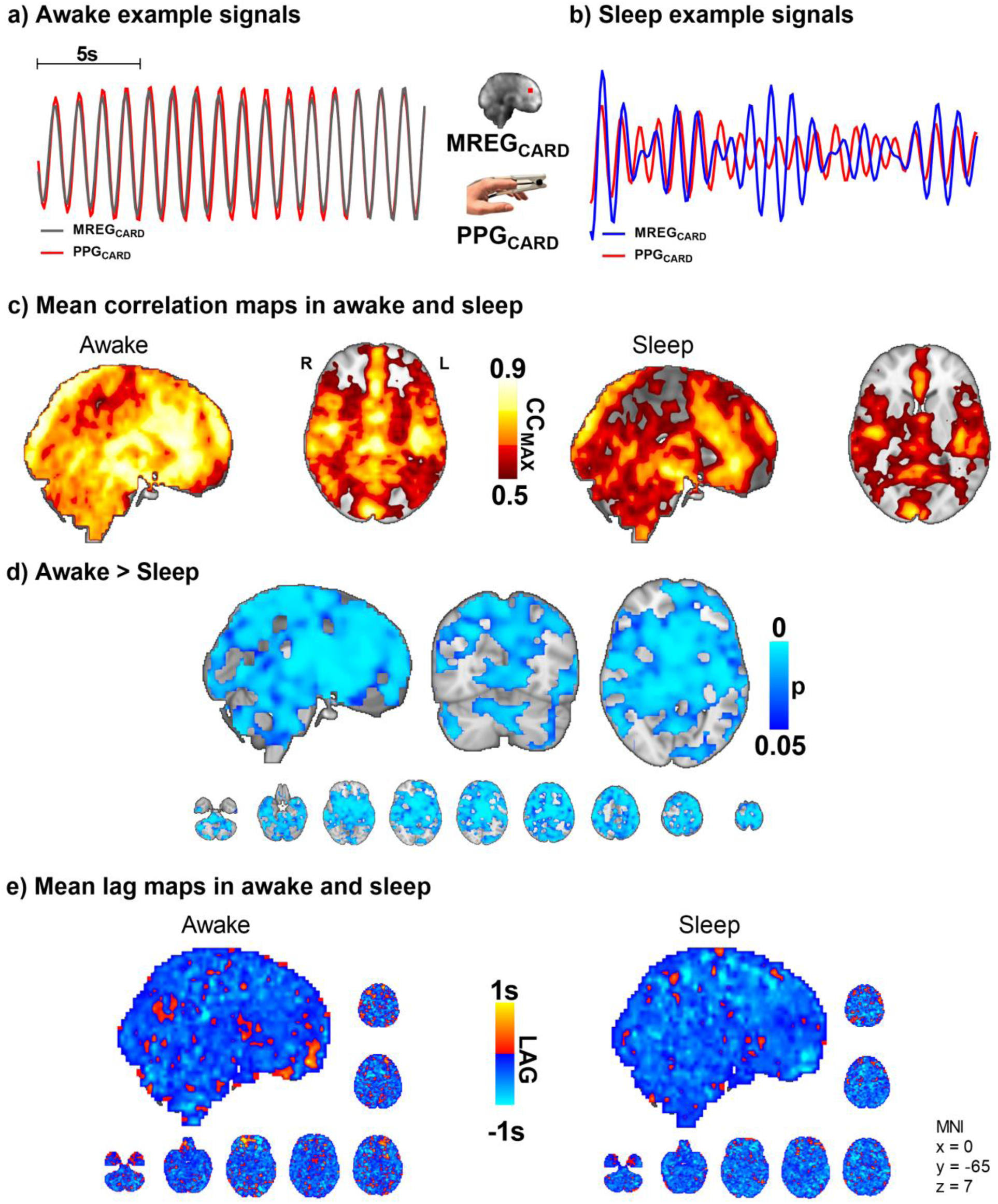
Sleep decreases the correlation between peripheral and brain cardiovascular measures of beat-to-beat pulses (n = 10). a-b) Representative fingertip and brain MREG cardiac signals during awake and sleep states. c-d) The group level maximum cross-correlation coefficients (CC_MAX_) between peripheral PPG_CARD_ and brain MREG_CARD_ during awake were nearly identical (CC_MAX_ ∼ 0.9) over most of the brain. Sleep reduced the beat-to-beat correlations significantly compared to awake state (p < 0.05), extending virtually over the whole brain. e) Lag between the signals indicated brain cardiovascular impulse preceding the peripheral as impulse propagates first into the brain. The lag did not change significantly during sleep.

Based on the above results, the cerebral vasoconstrictions may modulate cardiovascular pulsation, as reflected in the amplitude, more in sleep than while awake. Vasomotor contractions narrow blood vessels and increase local blood pressure^37^, which causes more rapid impulse propagation. The speed-up dephases the temporal synchrony between peripheral and brain cardiovascular signals (Fig. 3a-d). As the vasomotor tone relaxation does the opposite (slows down), the overall change in the averaged lag structure between the signals over the whole brain remains unchanged (Fig. 3e).

As there were strong vasomotor contractions in sleep both in peripheral and brain MREG_CARD_ signals, we proceeded to assess how the arterial vasomotor waves in the amplitude envelope correlated with the venous BOLD signal. We compared the regional brain vasomotor tone modulations with the downstream venous BOLD signal between awake and sleep states (Fig. 4). In sleep, the MREG_CHe_ and MREG_BOLD_ correlations were significantly higher in the visual cortex, cerebellum, and the parasagittal area, and in proximity to large veins (whole-brain analysis, p < 0.05, Fig. 4d). In other words, the cerebral arterial vasomotor waves and venous BOLD fluctuations become more synchronous during sleep in these regions, which were previously shown to present slow wave EEG activity reflecting increased fluid transport during sleep^12^. As before, the lag structure between these signals remained complex and unchanged by sleep (Fig. 4e). Over the whole brain the average on the BOLD signal preceded the CHe signal by 0.07 ± 1.4 seconds in waking, whereas in sleep the CHe precedes BOLD by 0.13 ± 1.4 seconds.

**Figure 4.**
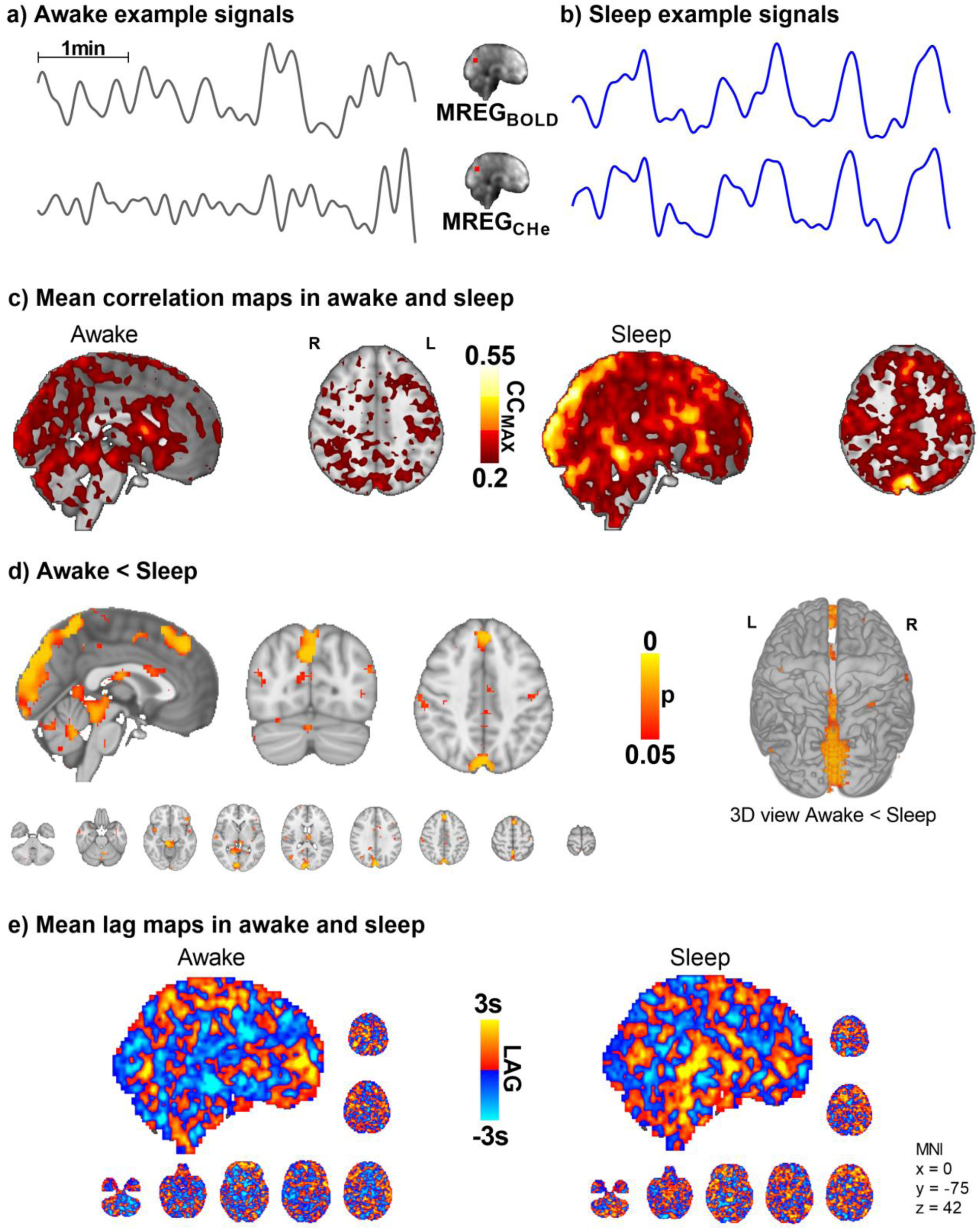
Intracranial synchrony of VLF modulations of arterial MREG_CHe_ pulsations and venous MREG_BOLD_ signals increases in sleep (n = 10). a-b) Example signals of MREG_CHe_, and MREG_BOLD_ during awake and sleep recordings. c-d) Cerebral signals between arterial pulsatility (MREG_CHe_) and vein-dominated baseline BOLD signal (MREG_BOLD_) harmonize during sleep, especially near large veins and the parasagittal area. e) There were no changes in the complex lag structure between the VLF changes in MREG_CHe_ vs. MREG_BOLD_, which was more heterogeneous than the cardiac pulsations presented in Fig 3.

### Peripheral blood volume vs. brain MREG_BOLD_ fluctuations

We next set out to examine the correlation between brain and peripheral blood volume pulsations by correlating the slow fingertip blood volume oscillations (PPG_VLF_) in periphery vs. brain MREG_BOLD_ VLF oscillations. The CC_MAX_ between PPG_VLF_ and MREG_BOLD_ was significantly higher in sleep compared to awake scans, extending widely over brain cortical regions (p < 0.05, Fig. 5). Interestingly, in the awake state, the fingertip blood volume oscillations follow the brain signal by a mean (± SD) of 0.03 ± 1.8 seconds, but in sleep, the peripheral oscillation precedes the brain MREG_BOLD_ signal by 2.1 ± 1.9 seconds. The phase difference was significant in the posterior cingulate gyrus, anterior default mode network area, and cerebellum in sleep (Fig. 5e).

**Figure 5.**
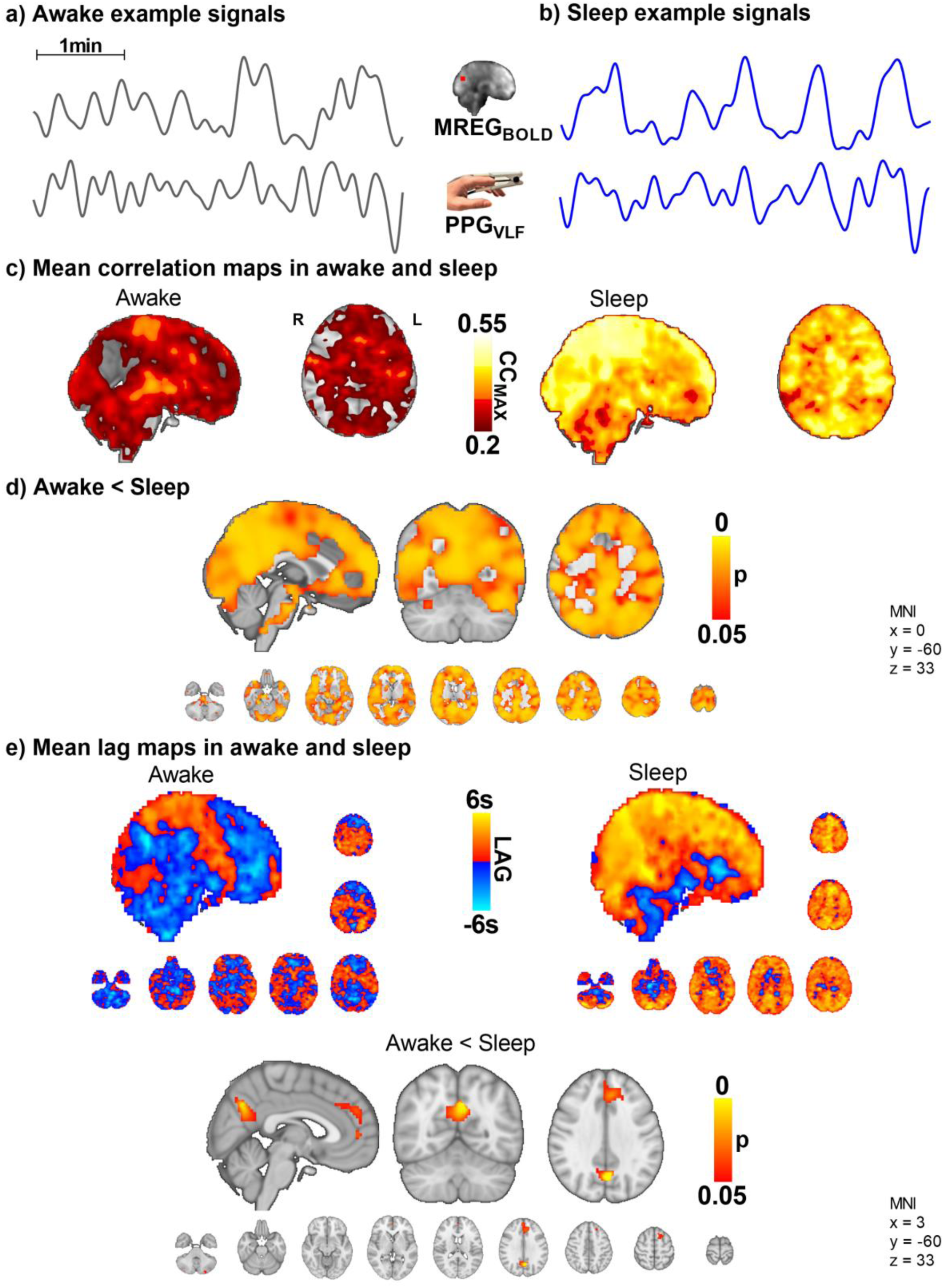
The VLF blood volume oscillations in the periphery (PPG_VLF_) and venous brain MREG_BOLD_ signal synchronize in sleep with a significant causal lag change in default mode brain areas (n = 10). a-b) Example signals of MREG_BOLD_ and PPG_VLF_ during awake and sleep states. c-d) The group level CC_MAX_ between MREG_BOLD_ and PPG_VLF_ during awake and sleep shows that correlations were significantly higher in sleep compared to awake (p < 0.05) widely in the brain. e) In sleep, the fingertip PPG_VLF_ signal has a significantly earlier onset than the MREG_BOLD_ signal (p < 0.05) in the posterior cingulate gyrus, the anterior default mode network area, and cerebellum.

### Peripheral arterial vasomotor CHe modulations vs. venous MREG_BOLD_ fluctuations

To ascertain how the peripheral arterial vasomotor tone PPG_CHe_ connects with venous brain MREG_BOLD_ fluctuations in sleep we compared these two signals. While awake, the peripheral vasomotor tone did not correlate with the brain venous BOLD signal, but during sleep the correlation between fingertip PPG_CHe_ and MREG_BOLD_ was significantly higher (p < 0.05, Fig. 6), extending over practically the whole brain. The change in the correlation between awake and sleep states was more significant than the correlation between MREG_BOLD_ vs PPG_VLF_ signals (Fig. 5d). On average, the peripheral PPG_CHe_ signal preceded brain MREG_BOLD_ signals by 0.7 ± 1.8 seconds in the awake state and 0.8 ± 1.6 seconds in sleep (Fig. 6e, non-significant).

**Figure 6.**
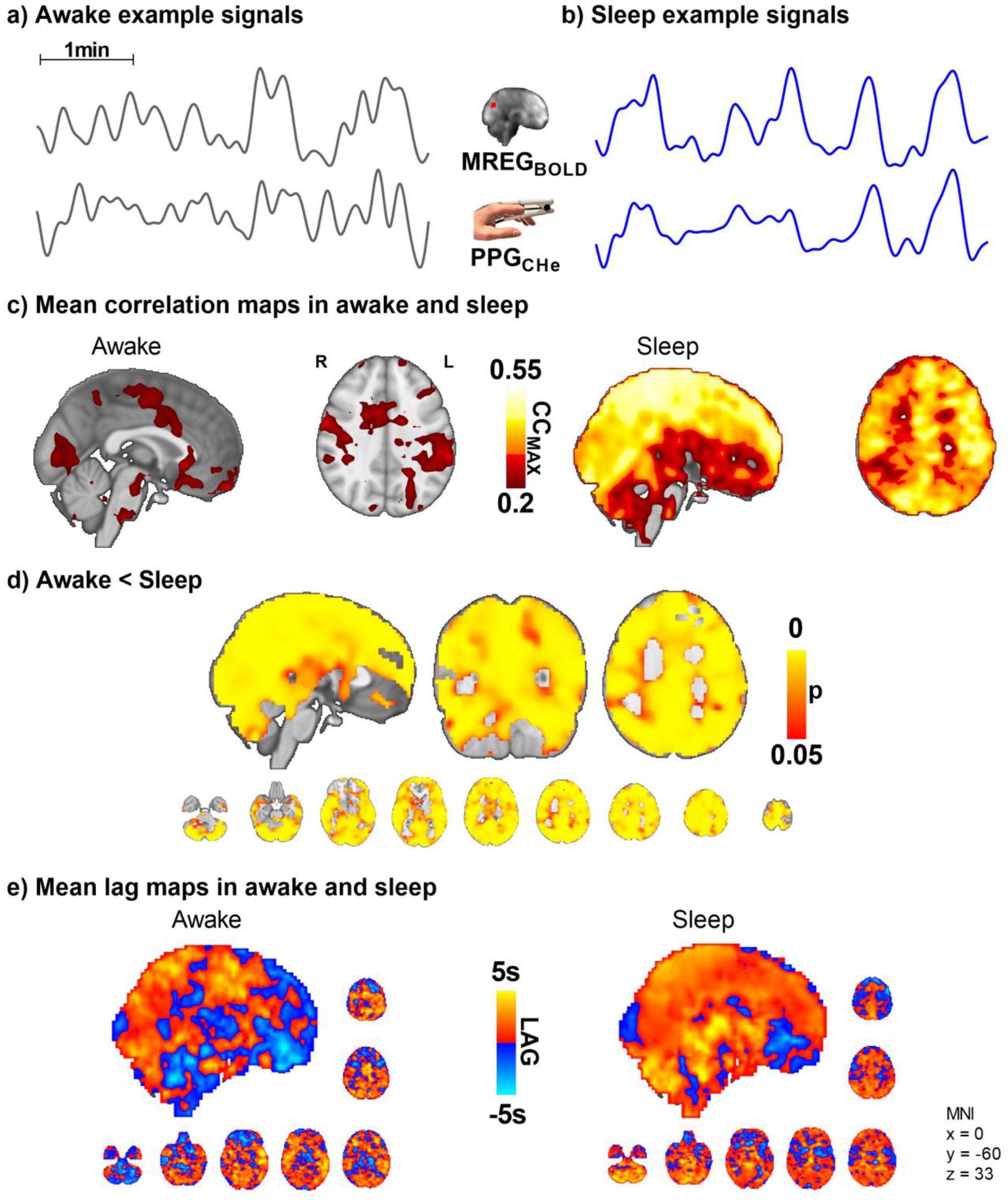
Peripheral vasomotor tone PPG_CHe_ and brain venous MREG_BOLD_ synchronize in sleep (n = 10). a-b) Representative MREG_BOLD_ and PPG_CHe_ signals during awake and sleep states. c-d) The correlation of the peripheral cardiovascular hemodynamic envelope (CHe) to the brain VLF MREG_BOLD_ signal shows that peripheral arterial vasomotor waves synchronize significantly with the cerebral BOLD signal during sleep. e) The lag shows a non-significant tendency towards increased lead in the peripheral and brain waves during sleep.

### Arterial PPG_CHe_ vs. brain arterial MREG_CHe_

We further examined the correlations between peripheral PPG_CHe_ and brain MREG_CHe_ (Fig. 7). The correlations between the CHe signals, which reflect amplitude modulations of the vasomotor tone, were highest in the arterial and sinus sagittal areas. There was no significant effect of sleep on synchrony. The brain signals preceded on average the PPG_CHe_ signal by 0.6 ± 1.4 seconds in awake state and by 0.03 ± 1.5 seconds in sleep, with a non-significant tendency for reversed lead from the peripheral signal. Examining the power of PPG_CHe_ and MREG_CHe_ did not reveal any statistically significant differences between awake and sleep.

**Figure 7.**
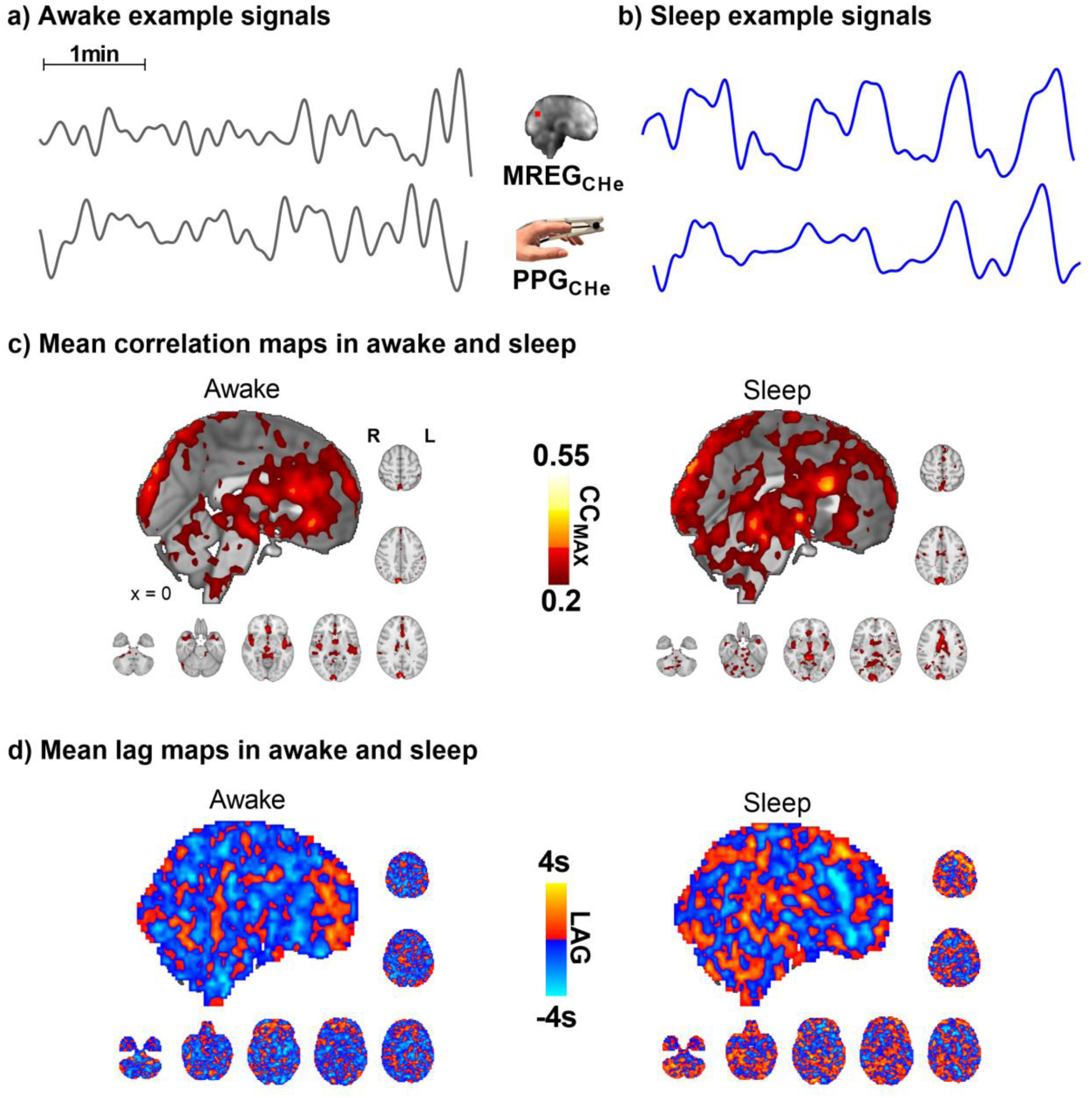
Correlation between cerebral and peripheral cardiac hemodynamic envelope signals (n = 10). a-b) Representative signals of MREG_CHe_ and PPG_CHe_ during awake and sleep states. c) The synchrony was highest in arterial and sinus sagittal areas, without significant changes between awake and sleep states. d) There are no significant changes in lag values between awake and sleep recordings.

## Discussion

In this study, we assessed the relationship between arterial cardiovascular pulsatility, vasomotor tone, and venous blood fluctuations in the brain and the periphery in awake and NREM sleep states. In sleep, the VLF waves between peripheral PPG and brain signals became significantly more synchronous, but the near-perfect cardiovascular impulse synchrony deteriorated. Within the brain, the arterial MREG_CHe_ vasomotor waves and venous MREG_BOLD_ signal fluctuations became more synchronized in parasagittal brain regions, where the power of VLF and cardiovascular signals also increased in the sleep state. Moreover, VLF (MREG_BOLD_) and cardiovascular (MREG_CARD_) powers in the brain were both significantly higher in sleep. In the periphery, however, the VLF power increased while the power of cardiovascular pulsations declined during sleep.

### Cardiovascular impulse synchrony between the periphery and brain vanishes in NREM sleep

The total power over the individual range of cardiac pulsation power increased in parietal areas, while the principal cardiac peak power was lower during sleep. In other words, the cardiovascular power widens over the entire frequency range in brain tissue during sleep, suggesting an increase in frequency modulation of the cardiovascular pulsation itself. The same phenomenon was also detected in peripheral PPG signals, reflecting the commonly known HRV, which is closely related to increased autonomic cardiovascular regulation in sleep^38^. Interestingly, however, our data show that the power of cardiovascular impulses changes in opposing directions in peripheral vs. brain signals (Fig. 2). This distinction may reflect a decreased peripheral oxygen consumption during sleep leads to a decrease in cardiac output, while cerebral blood flow increases in the brain^39^. Moreover, in the present study, the MREG power increase showed a somewhat wider spatial distribution along the sagittal sinus than in our previous analysis^12^, probably due to a more restricted cardiac power analysis centered around the principal cardiac peak power, instead of the full cardiac range of the whole group.

Also, we show for the first time that the high synchrony of the brain and peripheral cardiovascular beat-to-beat pulsatility seen in the awake state also disappears in NREM sleep, c.f. Fig. 3. Brief arousals, which are associated with sympathetic nervous system activation and consequent vasomotor contraction, are accompanied by substantial, transient elevations in blood pressure and HR^40–42^. Furthermore, the high blood pressure or vasomotor tone contractions also increase the speed of impulse propagation^37,43^, which is then visible as phase desynchronization between peripheral and brain pulsations (Fig. 3a-b). During sleep, the cardiovascular signals showed marked vasomotor contractions, as indicated by cardiac impulse amplitude drops, and reflected as modulation of the amplitude envelope, i.e. MREG_CHe_, figures 3-5. Since the coherence of the cardiovascular impulse amplitudes in brain MREG_CHe_ vs. peripheral PPG_CHe_ (Fig. 7) did not significantly differ between sleep and awake states, the desynchronization of the time domain signals must stem from signal dephasing. Thus, the vasomotor tone changes may be stronger in the brain than in the peripheral tissue. As the correlation values of the peripheral PPG (arterial tone waves and blood volume, Fig. 5-6) and brain slow waves reached a modest correlation coefficient of 0.55, and nearly perfect synchrony with a correlation of 0.9 in awake cardiovascular impulses (Fig. 3), there may be currently unidentified sources of modulation. An intriguing option for such modulatory interaction is the default mode brain network which is known to reflect changes in consciousness^44^. Here the default mode network BOLD signal started to follow the peripheral vasomotor CHe waves by some 2.1 seconds (Fig. 5e) in sleep and thus, the default mode seems to be connected to the modulation of the increased vasomotor waves of the body.

### Arterial vasomotor wave - CHe

The detection of cardiovascular signals with MREG has been previously shown to be very accurate; standard physiological measurement methods have shown virtually one-to-one correlation between MREG vs. PPG in multiple datasets^28,45^. Also in this study, the correlation between PPG and MREG was again 0.9 in arterial areas of the resting state awake scan, strongly indicating that individual cardiac filtered MREG signals reflect the arterial physiology especially close to the cerebral arteries. The envelope laid over the individual cardiovascular peaks reflects the amplitude changes of successive cardiovascular pulses locally. In terms of MR signal physics, each arterial impulse induces a fast drop in T2* weighted MREG signal due to a sudden pressure-induced disturbance in water spin precession- and phase^4,5^. In the veins, however, the dominating signal source has shown to be the *slow* susceptibility-weighted signal where the change in the venous Hb/HbO_2_ ratio increases T2* weighted signal 3-5 sec after the activation-induced vasodilation^6,7^. As the arteries vasodilate first and the venous signal follows, these both effects can be separated using the fast scanning technique, such as the 10 Hz MREG^8,9^. MREG indicated a 1.3 sec time difference of the arterial dilation vs. venous BOLD signal elevation after cued visual brain activation^8^.

We wanted to investigate how the hemodynamic coupling between the intracranial arterial MREG_CHe_ and venous MREG_BOLD_ signal differs between awake and sleep states. The synchrony between the arterial MREG_CHe_ and venous MREG_BOLD_ signal seen in the awake recordings was significantly higher in parasagittal areas during NREM sleep (Fig. 4). Around these parietal parasagittal areas, the powers both of cardiovascular impulses MREG_CARD_ and MREG_BOLD_ venous waves were higher in sleep, c.f. Fig. 2. These brain areas have increased slow delta EEG power during sleep, predicting increased ISF clearance^12,46^. Moreover, intrathecal gadolinium MRI contrast agent and protein solutes accumulated in the very same parasagittal areas in human glymphatic studies^47,48^.

Based on the spatial overlaps, we believe that the increased synchrony between the arterial MREG_CHe_ and venous MREG_BOLD_ during sleep reflects increased convection between the two compartments, a phenomenon detected using these two MREG contrasts. This finding could reflect the increased power of the pulsations in these brain areas. The increased synchrony between the arterial and venous pulsations could suggest that a local mechanism allows pulsations to pass more readily from the arterial to venous side, either via the glia limitans of the blood-brain barrier as proposed by the mechanism of water transport and/or via increased capillary blood flow and recruitment. Addressing this issue requires more detailed investigations utilizing microscopic methods and tools specifically tailored to detect water exchange and blood-brain barrier permeability.

### Venous BOLD fluctuations

Human sleep manifests in strong increases in the VLF < 0.1 Hz BOLD signal fluctuations^49–52^. In the awake state, the functional BOLD signal exhibits robust coupling with neuronal activity^53–57^. However, in sleep^12,49,50,58^, or anesthesia^44,59,60^, this local coupling seems to be dominated by slowly propagating 0.03-0.04 Hz waves^2,61–63^. While some researchers argue that neurovascular coupling is maintained in sleep^52^, others showed that a 0.04 Hz vasomotor wave appears under anesthesia, which masks the signals that were functionally coupled to neuronal activity^59^. Consistent with these findings, a recent study presented compelling evidence of the simultaneous influence of these two signaling mechanisms on BOLD signals in the human brain. One mechanism propagates into a vasomotor wave type, while the other establishes functional connectivity with a standing wave type coupled to neuronal activity^64^.

A recent study on mice indicates that innervations from the locus coeruleus produce slow noradrenaline level waves that drive widespread vasomotor waves at around 0.03 Hz to control sleep state architecture^65^. Human data show that, in a state of reduced consciousness, the brain exhibits slow co-activation patterns induced by the cholinergic nucleus basalis of Meynert^51^, whereas an invasive study in monkeys showed that detection of this coactivation in fMRI data is encumbered by physiological pulsations^66^. Those studies align with fast MREG measurements showing an increase in slow VLF changes in sleep, where the respiratory and cardiac pulsations similarly increased according to a separate mechanism during sleep^12^.

Waves involved in the regulation of blood pressure in the ∼0.1 Hz range may explain much of the BOLD contrast signal in awake humans^67,68^. The present study shows that during NREM sleep, the slower variations appear both in the global BOLD signal and the peripheral signal, especially in the < 0.05 Hz frequency range (Fig. 2b). The increase in power in the peripheral vasomotor power occurs in the same frequency ranges as were seen globally in the brain. In both regions, strongly dominant waves appear in the ∼ 0.02 and 0.04 Hz ranges. It has been shown that spectral analysis of the mean flow velocity of the middle cerebral artery increases in power at 0.02 Hz during NREM sleep^69^. Sleep also increases in power at 0.05 Hz^52^. In the VLF range, there are individual frequency peaks similar to the cardiac and respiration ranges. It should be noted that cardiorespiratory aliasing occurs significantly in the VLF range when TR is more than 100 ms^3^. Because of this, we suggest that the VLF power was measured more accurately in this paper. Indeed, we propose that sleeping hemodynamic waves appear to be associated with vasomotion and that < 0.1 Hz vasomotor waves may explain much of the BOLD signal during NREM N1-N2 sleep; we infer that this may be a significant drive of fluid transport. Furthermore, as the vasomotor wave intensifies, there is an associated acceleration of tracer movement along the perivascular space^18^.

Results of this study show that in NREM sleep, part of the VLF signal indeed originates in a systemic-origin vasomotor wave. Furthermore, the increase in the vasomotor wave power and synchrony appearing in the human brain in NREM sleep may significantly affect the increased brain fluid transport in sleep. The default mode network seems to start following the peripheral vasomotor waves supporting earlier evidence on the modulatory role of it in changes of consciousness. In addition, the venous signal harmonizes with the arterial signal in parasagittal brain areas during sleep, which may also enable CSF flow. Finally, we find that cardiac pulsatility is not coherent with peripheral pulsatility during NREM sleep. With the advent of fast fMRI techniques, it is now possible to visualize the drivers of CSF flow. This study provides additional information on how cardiovascular changes occurring in NREM sleep affect brain pulsations, which implies that aspects of cerebral circulation could play a significant role in driving brain fluid transport.

### Limitations

25 subjects participated in the study^12^, but complete data including sleep scores, MREG, and PPG data for both awake and sleep scans were available for only 11. Exclusion criteria primarily resulted from continuous artifacts in EEG, and a detailed account is provided in Helakari et al. (2022)^12^. Data from one subject were also excluded because of excessive noise in the PPG signal, resulting in a total number of ten subjects.

Our objective was to investigate the similarity of signals at the same phase in awake and NREM sleep states. Because we wanted to study similarity and phase locking, we ruled out consideration of a negative correlation, which may be a matter for future investigation. Also, since the cardiac power increase during sleep was smaller compared to the VLF and respiratory pulsations changes, it could be proposed that the modulating factor affecting both CHe and BOLD signals could be the respiratory pulsations, which are known to modulate the cardiovascular signal, whereas the veins mediate the respiratory effects^9,12,70^. In this regard, it is noteworthy that the peripheral arterial amplitude vasomotion (PPG_CHe_) and volume (PPG_VLF_) fluctuations and the venous MREG_BOLD_ signal become markedly more synchronous in sleep than with the intracranial arterial amplitude MREG_CHe_. The venous BOLD lag also increased significantly compared to peripheral blood volume PPG_VLF_ oscillations, Fig. 5. One might expect that the intracranial arterial CHe modulations should be more in synchrony with the venous BOLD signal than with peripheral signals, but the intracranial synchrony seen in the present study appear to be confined to areas showing sleep-related increases in solute transport^47^, c.f. Fig. 4. Thus, the synchrony of local and peripheral influences must be considered together.

The awake scan was measured at 4:00-6:00 P.M. and the sleep scan at 6:00-8:00 A.M. Circadian rhythms may have confounded our results but to minimize that possibility, we selected for analysis those continuous 5-minute segments that included the highest amount of NREM N2 sleep according to AASM sleep scoring criteria. We also used 24-hour sleep deprivation to increase sleep pressure and help subjects fall asleep in the scanner^71,72^.

## Acknowledgments

We would like to thank all study subjects for their participation in the study. We also thank Jani Häkli, Tarja Holtinkoski, Aleksi Rasila, Taneli Hautaniemi, Miia Lampinen, and others who assisted in measurements or participated otherwise. We are thankful for the devices and data provided by Oura Health. We wish to acknowledge Jussi Kantola for data management and reconstruction of MREG data and the CSC – IT Center for Science Ltd., Finland for generous computational resources, and Prof. Paul Cumming for comments and proofreading on the manuscript.

## Conflict of interest statement

None declared.

## Notes

### Competing Interest Statement

The authors have declared no competing interest.

## References

1. Mosso A. Sulla Circolazione del Sangue nel Cervello Dell’Uomo: Ricerche Sfigmografiche. Roma: Coi tipi del Salviucci. Published online 1880.

2. Kiviniemi V, Wang X, Korhonen V, et al. Ultra-fast magnetic resonance encephalography of physiological brain activity - Glymphatic pulsation mechanisms? J Cereb Blood Flow Metab. 2016;36(6):1033–1045. doi:10.1177/0271678X15622047

3. Huotari N, Raitamaa L, Helakari H, et al. Sampling Rate Effects on Resting State fMRI Metrics. Front Neurosci. 2019;13. doi:10.3389/fnins.2019.00279

4. von Schulthess GK, Higgins CB. Blood flow imaging with MR: spin-phase phenomena. Radiology. 1985;157(3):687–695. doi:10.1148/radiology.157.3.2997836

5. Duyn JH. Steady state effects in fast gradient echo magnetic resonance imaging. Magn Reson Med. 1997;37(4):559–568. doi:10.1002/mrm.1910370414

6. Ogawa S, Lee TM. Magnetic resonance imaging of blood vessels at high fields:In vivo andin vitro measurements and image simulation. Magn Reson Med. 1990;16(1):9–18. doi:10.1002/mrm.1910160103

7. Buxton RB. Dynamic models of BOLD contrast. Neuroimage. 2012;62(2):953–961. doi:10.1016/j.neuroimage.2012.01.012

8. Huotari N, Tuunanen J, Raitamaa L, et al. Cardiovascular Pulsatility Increases in Visual Cortex Before Blood Oxygen Level Dependent Response During Stimulus. Front Neurosci. 2022;16. doi:10.3389/fnins.2022.836378

9. Raitamaa L, Korhonen V, Huotari N, et al. Breath hold effect on cardiovascular brain pulsations – A multimodal magnetic resonance encephalography study. Journal of Cerebral Blood Flow & Metabolism. 2019;39(12):2471–2485. doi:10.1177/0271678X18798441

10. Özbay PS, Chang C, Picchioni D, et al. Sympathetic activity contributes to the fMRI signal. Commun Biol. 2019;2(1):421. doi:10.1038/s42003-019-0659-0

11. Tong Y, Frederick B deB. Concurrent fNIRS and fMRI processing allows independent visualization of the propagation of pressure waves and bulk blood flow in the cerebral vasculature. Neuroimage. 2012;61(4):1419–1427. doi:10.1016/j.neuroimage.2012.03.009

12. Helakari H, Korhonen V, Holst SC, et al. Human NREM Sleep Promotes Brain-Wide Vasomotor and Respiratory Pulsations. The Journal of Neuroscience. 2022;42(12):2503–2515. doi:10.1523/JNEUROSCI.0934-21.2022

13. Iliff JJ, Wang M, Liao Y, et al. A Paravascular Pathway Facilitates CSF Flow Through the Brain Parenchyma and the Clearance of Interstitial Solutes, Including Amyloid β. Sci Transl Med. 2012;4(147). doi:10.1126/scitranslmed.3003748

14. Mestre H, Tithof J, Du T, et al. Flow of cerebrospinal fluid is driven by arterial pulsations and is reduced in hypertension. Nat Commun. 2018;9(1):4878. doi:10.1038/s41467-018-07318-3

15. Meng Y, Abrahao A, Heyn CC, et al. Glymphatics Visualization after Focused Ultrasound-Induced Blood–Brain Barrier Opening in Humans. Ann Neurol. 2019;86(6):975–980. doi:10.1002/ana.25604

16. Nedergaard M. Garbage Truck of the Brain. Science (1979). 2013;340(6140):1529–1530. doi:10.1126/science.1240514

17. Santisakultarm TP, Cornelius NR, Nishimura N, et al. In vivo two-photon excited fluorescence microscopy reveals cardiac- and respiration-dependent pulsatile blood flow in cortical blood vessels in mice. American Journal of Physiology-Heart and Circulatory Physiology. 2012;302(7):H1367–H1377. doi:10.1152/ajpheart.00417.2011

18. van Veluw SJ, Hou SS, Calvo-Rodriguez M, et al. Vasomotion as a Driving Force for Paravascular Clearance in the Awake Mouse Brain. Neuron. 2020;105(3):549–561.e5. doi:10.1016/j.neuron.2019.10.033

19. Aldea R, Weller RO, Wilcock DM, Carare RO, Richardson G. Cerebrovascular Smooth Muscle Cells as the Drivers of Intramural Periarterial Drainage of the Brain. Front Aging Neurosci. 2019;11. doi:10.3389/fnagi.2019.00001

20. Assländer J, Zahneisen B, Hugger T, et al. Single shot whole brain imaging using spherical stack of spirals trajectories. Neuroimage. 2013;73:59–70. doi:10.1016/j.neuroimage.2013.01.065

21. Hugger T, Zahneisen B, LeVan P, et al. Fast Undersampled Functional Magnetic Resonance Imaging Using Nonlinear Regularized Parallel Image Reconstruction. PLoS One. 2011;6(12):e28822. doi:10.1371/journal.pone.0028822

22. Korhonen V, Hiltunen T, Myllylä T, et al. Synchronous Multiscale Neuroimaging Environment for Critically Sampled Physiological Analysis of Brain Function: Hepta-Scan Concept. Brain Connect. 2014;4(9):677–689. doi:10.1089/brain.2014.0258

23. Jenkinson M, Beckmann CF, Behrens TEJ, Woolrich MW, Smith SM. FSL. Neuroimage. 2012;62(2):782–790. doi:10.1016/j.neuroimage.2011.09.015

24. Smith SM. Fast robust automated brain extraction. Hum Brain Mapp. 2002;17(3):143–155. doi:10.1002/hbm.10062

25. Cox RW. AFNI: Software for Analysis and Visualization of Functional Magnetic Resonance Neuroimages. Computers and Biomedical Research. 1996;29(3):162–173. doi:10.1006/cbmr.1996.0014

26. Jenkinson M, Bannister P, Brady M, Smith S. Improved Optimization for the Robust and Accurate Linear Registration and Motion Correction of Brain Images. Neuroimage. 2002;17(2):825–841. doi:10.1016/S1053-8119(02)91132-8

27. Kananen J, Järvelä M, Korhonen V, et al. Increased interictal synchronicity of respiratory related brain pulsations in epilepsy. Journal of Cerebral Blood Flow & Metabolism. 2022;42(10):1840–1853. doi:10.1177/0271678X221099703

28. Järvelä M, Kananen J, Korhonen V, Huotari N, Ansakorpi H, Kiviniemi V. Increased very low frequency pulsations and decreased cardiorespiratory pulsations suggest altered brain clearance in narcolepsy. Communications Medicine. 2022;2(1):122. doi:10.1038/s43856-022-00187-4

29. Shaffer F, McCraty R, Zerr CL. A healthy heart is not a metronome: an integrative review of the heart’s anatomy and heart rate variability. Front Psychol. 2014;5. doi:10.3389/fpsyg.2014.01040

30. Gasparini F, Grossi A, Giltri M, Bandini S. Personalized PPG Normalization Based on Subject Heartbeat in Resting State Condition. Signals. 2022;3(2):249–265. doi:10.3390/signals3020016

31. Schreiber SJ, Franke U, Doepp F, Staccioli E, Uludag K, Valdueza JM. Dopplersonographic measurement of global cerebral circulation time using echo contrast-enhanced ultrasound in normal individuals and patients with arteriovenous malformations. Ultrasound Med Biol. 2002;28(4):453–458. doi:10.1016/S0301-5629(02)00477-5

32. Allen J. Photoplethysmography and its application in clinical physiological measurement. Physiol Meas. 2007;28(3):R1–R39. doi:10.1088/0967-3334/28/3/R01

33. Abay TY, Kyriacou PA. Photoplethysmography for blood volumes and oxygenation changes during intermittent vascular occlusions. J Clin Monit Comput. 2018;32(3):447–455. doi:10.1007/s10877-017-0030-2

34. Allen PJ, Josephs O, Turner R. A Method for Removing Imaging Artifact from Continuous EEG Recorded during Functional MRI. Neuroimage. 2000;12(2):230–239. doi:10.1006/nimg.2000.0599

35. Allen PJ, Polizzi G, Krakow K, Fish DR, Lemieux L. Identification of EEG Events in the MR Scanner: The Problem of Pulse Artifact and a Method for Its Subtraction. Neuroimage. 1998;8(3):229–239. doi:10.1006/nimg.1998.0361

36. Lohmann G, Stelzer J, Lacosse E, et al. LISA improves statistical analysis for fMRI. Nat Commun. 2018;9(1):4014. doi:10.1038/s41467-018-06304-z

37. Myllylä TS, Elseoud AA, Sorvoja HSS, et al. Fibre optic sensor for non-invasive monitoring of blood pressure during MRI scanning. J Biophotonics. 2011;4(1-2):98–107. doi:10.1002/jbio.200900105

38. Mendez MO, Matteucci M, Castronovo V, Strambi LF, Cerutti S, Bianchi AM. Sleep staging from Heart Rate Variability: time-varying spectral features and Hidden Markov Models. Int J Biomed Eng Technol. 2010;3(3/4):246. doi:10.1504/IJBET.2010.032695

39. Coote JH. Respiratory and Circulatory Control During Sleep. Journal of Experimental Biology. 1982;100(1):223–244. doi:10.1242/jeb.100.1.223

40. Burgess HJ, Kleiman J, Trinder J. Cardiac activity during sleep onset. Psychophysiology. 1999;36(3):S0048577299980198. doi:10.1017/S0048577299980198

41. Liu X, Duyn JH. Time-varying functional network information extracted from brief instances of spontaneous brain activity. Proceedings of the National Academy of Sciences. 2013;110(11):4392–4397. doi:10.1073/pnas.1216856110

42. Carrington MJ, Barbieri R, Colrain IM, Crowley KE, Kim Y, Trinder J. Changes in cardiovascular function during the sleep onset period in young adults. J Appl Physiol. 2005;98(2):468–476. doi:10.1152/japplphysiol.00702.2004

43. Finnegan E, Davidson S, Harford M, et al. Pulse arrival time as a surrogate of blood pressure. Sci Rep. 2021;11(1):22767. doi:10.1038/s41598-021-01358-4

44. Kiviniemi V, Haanpää H, Kantola JH, et al. Midazolam sedation increases fluctuation and synchrony of the resting brain BOLD signal. Magn Reson Imaging. 2005;23(4):531–537. doi:10.1016/j.mri.2005.02.009

45. Tuovinen T, Kananen J, Rajna Z, et al. The variability of functional MRI brain signal increases in Alzheimer’s disease at cardiorespiratory frequencies. Sci Rep. 2020;10(1):21559. doi:10.1038/s41598-020-77984-1

46. Ding F, O’Donnell J, Xu Q, Kang N, Goldman N, Nedergaard M. Changes in the composition of brain interstitial ions control the sleep-wake cycle. Science (1979). 2016;352(6285):550–555. doi:10.1126/science.aad4821

47. Ringstad G, Eide PK. Cerebrospinal fluid tracer efflux to parasagittal dura in humans. Nat Commun. 2020;11(1):354. doi:10.1038/s41467-019-14195-x

48. Albayram MS, Smith G, Tufan F, et al. Non-invasive MR imaging of human brain lymphatic networks with connections to cervical lymph nodes. Nat Commun. 2022;13(1):203. doi:10.1038/s41467-021-27887-0

49. Fukunaga M, Horovitz SG, van Gelderen P, et al. Large-amplitude, spatially correlated fluctuations in BOLD fMRI signals during extended rest and early sleep stages. Magn Reson Imaging. 2006;24(8):979–992. doi:10.1016/j.mri.2006.04.018

50. Horovitz SG, Fukunaga M, de Zwart JA, et al. Low frequency BOLD fluctuations during resting wakefulness and light sleep: A simultaneous EEG-fMRI study. Hum Brain Mapp. 2008;29(6):671–682. doi:10.1002/hbm.20428

51. Liu X, de Zwart JA, Schölvinck ML, et al. Subcortical evidence for a contribution of arousal to fMRI studies of brain activity. Nat Commun. 2018;9(1):395. doi:10.1038/s41467-017-02815-3

52. Fultz NE, Bonmassar G, Setsompop K, et al. Coupled electrophysiological, hemodynamic, and cerebrospinal fluid oscillations in human sleep. Science (1979). 2019;366(6465):628–631. doi:10.1126/science.aax5440

53. Ogawa S, Tank DW, Menon R, et al. Intrinsic signal changes accompanying sensory stimulation: functional brain mapping with magnetic resonance imaging. Proceedings of the National Academy of Sciences. 1992;89(13):5951–5955. doi:10.1073/pnas.89.13.5951

54. Biswal B, Zerrin Yetkin F, Haughton VM, Hyde JS. Functional connectivity in the motor cortex of resting human brain using echo-planar mri. Magn Reson Med. 1995;34(4):537–541. doi:10.1002/mrm.1910340409

55. Drew PJ, Duyn JH, Golanov E, Kleinfeld D. Finding coherence in spontaneous oscillations. Nat Neurosci. 2008;11(9):991–993. doi:10.1038/nn0908-991

56. Rayshubskiy A, Wojtasiewicz TJ, Mikell CB, et al. Direct, intraoperative observation of ∼0.1Hz hemodynamic oscillations in awake human cortex: Implications for fMRI. Neuroimage. 2014;87:323–331. doi:10.1016/j.neuroimage.2013.10.044

57. Fox MD, Snyder AZ, Vincent JL, Corbetta M, Van Essen DC, Raichle ME. The human brain is intrinsically organized into dynamic, anticorrelated functional networks. Proceedings of the National Academy of Sciences. 2005;102(27):9673–9678. doi:10.1073/pnas.0504136102

58. Picchioni D, Özbay PS, Mandelkow H, et al. Autonomic arousals contribute to brain fluid pulsations during sleep. Neuroimage. 2022;249:118888. doi:10.1016/j.neuroimage.2022.118888

59. Ma Y, Shaik MA, Kozberg MG, et al. Resting-state hemodynamics are spatiotemporally coupled to synchronized and symmetric neural activity in excitatory neurons. Proceedings of the National Academy of Sciences. 2016;113(52). doi:10.1073/pnas.1525369113

60. Kiviniemi V, Jauhiainen J, Tervonen O, et al. Slow vasomotor fluctuation in fMRI of anesthetized child brain. Magn Reson Med. 2000;44(3):373–378. doi:10.1002/1522-2594(200009)44:3<373::AID-MRM5>3.0.CO;2-P

61. Gu Y, Sainburg LE, Kuang S, et al. Brain Activity Fluctuations Propagate as Waves Traversing the Cortical Hierarchy. Cerebral Cortex. 2021;31(9):3986–4005. doi:10.1093/cercor/bhab064

62. Raut R V., Snyder AZ, Mitra A, et al. Global waves synchronize the brain’s functional systems with fluctuating arousal. Sci Adv. 2021;7(30). doi:10.1126/sciadv.abf2709

63. Yousefi B, Shin J, Schumacher EH, Keilholz SD. Quasi-periodic patterns of intrinsic brain activity in individuals and their relationship to global signal. Neuroimage. 2018;167:297–308. doi:10.1016/j.neuroimage.2017.11.043

64. Bolt T, Nomi JS, Bzdok D, et al. A parsimonious description of global functional brain organization in three spatiotemporal patterns. Nat Neurosci. 2022;25(8):1093–1103. doi:10.1038/s41593-022-01118-1

65. Kjaerby C, Andersen M, Hauglund N, et al. Memory-enhancing properties of sleep depend on the oscillatory amplitude of norepinephrine. Nat Neurosci. 2022;25(8):1059–1070. doi:10.1038/s41593-022-01102-9

66. Chang C, Leopold DA, Schölvinck ML, et al. Tracking brain arousal fluctuations with fMRI. Proceedings of the National Academy of Sciences. 2016;113(16):4518–4523. doi:10.1073/pnas.1520613113

67. Whittaker JR, Driver ID, Venzi M, Bright MG, Murphy K. Cerebral Autoregulation Evidenced by Synchronized Low Frequency Oscillations in Blood Pressure and Resting-State fMRI. Front Neurosci. 2019;13. doi:10.3389/fnins.2019.00433

68. Attarpour A, Ward J, Chen JJ. Vascular origins of low-frequency oscillations in the cerebrospinal fluid signal in resting-state fMRI: Interpretation using photoplethysmography. Hum Brain Mapp. 2021;42(8):2606–2622. doi:10.1002/hbm.25392

69. Lee WJ, Jung KH, Park HM, et al. Periodicity of cerebral flow velocity during sleep and its association with white-matter hyperintensity volume. Sci Rep. 2019;9(1):15510. doi:10.1038/s41598-019-52029-4

70. Raitamaa L, Huotari N, Korhonen V, et al. Spectral analysis of physiological brain pulsations affecting the BOLD signal. Hum Brain Mapp. 2021;42(13):4298–4313. doi:10.1002/hbm.25547

71. Portas CM, Krakow K, Allen P, Josephs O, Armony JL, Frith CD. Auditory Processing across the Sleep-Wake Cycle. Neuron. 2000;28(3):991–999. doi:10.1016/S0896-6273(00)00169-0

72. Horovitz SG, Braun AR, Carr WS, et al. Decoupling of the brain’s default mode network during deep sleep. Proceedings of the National Academy of Sciences. 2009;106(27):11376–11381. doi:10.1073/pnas.0901435106

